# ULTRAFAST STRUCTURAL CHANGES DIRECT THE FIRST MOLECULAR EVENTS OF VISION

**DOI:** 10.1101/2022.10.14.511948

**Authors:** Thomas Gruhl, Tobias Weinert, Matthew Rodrigues, Christopher J Milne, Giorgia Ortolani, Karol Nass, Eriko Nango, Saumik Sen, Philip J M Johnson, Claudio Cirelli, Antonia Furrer, Sandra Mous, Petr Skopintsev, Daniel James, Florian Dworkowski, Petra Båth, Demet Kekilli, Dmitry Ozerov, Rie Tanaka, Hannah Glover, Camila Bacellar, Steffen Brünle, Cecilia M Casadei, Azeglio D Diethelm, Dardan Gashi, Guillaume Gotthard, Ramon Guixà-González, Yasumasa Joti, Victoria Kabanova, Gregor Knopp, Elena Lesca, Pikyee Ma, Isabelle Martiel, Jonas Mühle, Shigeki Owada, Filip Pamula, Daniel Sarabi, Oliver Tejero, Ching-Ju Tsai, Niranjan Varma, Anna Wach, Sébastien Boutet, Kensuke Tono, Przemyslaw Nogly, Xavier Deupi, So Iwata, Richard Neutze, Jörg Standfuss, Gebhard FX Schertler, Valerie Panneels

**Author notes:** equal contribution (Tobias Weinert, Matthew Rodrigues). Christopher Milne: European XFEL GmbH, 22869 Schenefeld, Germany; Antonia Furrer: Biologics Center, Novartis Institutes for Biomedical Research, Novartis Campus, Basel, Switzerland; Sandra Mous : Linac Coherent Light Source, SLAC National Accelerator Laboratory, Menlo Park, CA 94025, USA; Petr Skopintsev: California Institute for Quantitative Biosciences (QB3), University of California, Berkeley, CA, USA; Daniel James: Department of Physics, Utah Valley University, Orem, UT 84058, USA; Steffen Brünle : Leiden Institute of Chemistry, Leiden University, Leiden, The Netherlands; Filip Pamula: Department of Molecular Biology and Genetics, Aarhus University, Denmark; Przemyslaw Nogly: Dioscuri Center For Structural Dynamics of Receptors, Faculty of Biochemistry, Biophysics and Biotechnology, Jagiellonian University in Kraków, 30-387 Kraków, Poland.

## Abstract

Vision is initiated by the rhodopsin family of light-sensitive G protein-coupled receptors (GPCRs). A photon is absorbed by the 11-*cis* retinal chromophore of rhodopsin which isomerises within 200 femtoseconds to the all-*trans* conformation, thereby initiating the cellular signal transduction processes that ultimately lead to vision. However, the intramolecular mechanism by which the photoactivated retinal induces the activation events inside rhodopsin remains elusive. In this work, we use ultrafast time-resolved crystallography at room temperature to determine how an isomerised twisted *all-trans* retinal stores the photon energy required to initiate protein conformational changes associated with the formation of the G protein-binding signalling state. The distorted retinal at 1 ps time-delay of photoactivation has pulled away from half of its numerous interactions with its binding pocket, and the excess of the photon energy is released through an anisotropic protein breathing motion in the direction of the extracellular space. Strikingly, the very early structural motions in the protein side chains of rhodopsin appear in regions involved in later stages of the conserved Class A GPCR activation mechanism. Our work sheds light on the earliest stages of vision in vertebrates and points to fundamental aspects of the molecular mechanisms of agonist-mediated GPCR activation.

---

Rhodopsin, the vertebrate receptor for low-light vision, is concentrated within the disk membranes of rod cells in the retina. Rhodopsin transforms the absorption of light into a physiological signal through conformational changes that activate the intracellular G protein transducin –a member of the Gi/o/t family– initiating a signalling cascade, resulting in electrical impulses sent to the brain and ultimately leading to visual perception. The structure of rhodopsin consists of seven transmembrane (TM) α-helices with an 11-*cis* retinal chromophore covalently bound through a protonated Schiff base (PSB) to Lys296^<u>7.43</u>^ of TM7 (underlined superscripts denote the general residue number for GPCRs^1^, see Methods). This buried ligand is located within the TM bundle towards the extracellular side, like in many Class A GPCRs. Retinal also contacts extracellular loop 2 (ECL2), which forms a lid over the chromophore and contains a highly conserved disulfide bridge (Cys110^<u>3.25</u>^-Cys187^<u>ECL2</u>^) connecting to the central helix TM3 (see Box 1 in reference^2^ for an overview of the activation microswitches in rhodopsin). Glu113^<u>3.28</u>^ provides a negatively charged counterion^3^ that forms a salt bridge with the PSB (**Fig. 1a-c**) and thereby participates in the stabilisation of this buried positive charge in the resting state of the receptor^4,5^. From our understanding of the evolution of visual pigments^6,7^ we know that originally, Glu181 (the “ancestral counterion”^8^, located in ECL2, which still plays the role of a complex counterion in invertebrates^6^) was the only residue able to neutralise the positive charge of the SB. In vertebrates, a second counterion Glu113 appeared during evolution and both residues are important for the activation mechanism. The PSB is still connected through a water-mediated hydrogen bond network to the ancestral counterion Glu181 (**Fig. 1c**). Structures of light-activated rhodopsin trapped at low-temperature^9,10^, of the late Meta II active state^11,12,13^ (bound to G proteins and G protein-mimicking peptides), and copious computational^14^, biochemical and spectroscopic studies have provided invaluable insights into the mechanism of signal transduction in rhodopsin (see^15^ for a review). However, methods that provide both a high spatial and temporal resolution are required to obtain a complete picture of the activation mechanism at the atomic scale from femtoseconds to milliseconds.

**Fig. 1.**
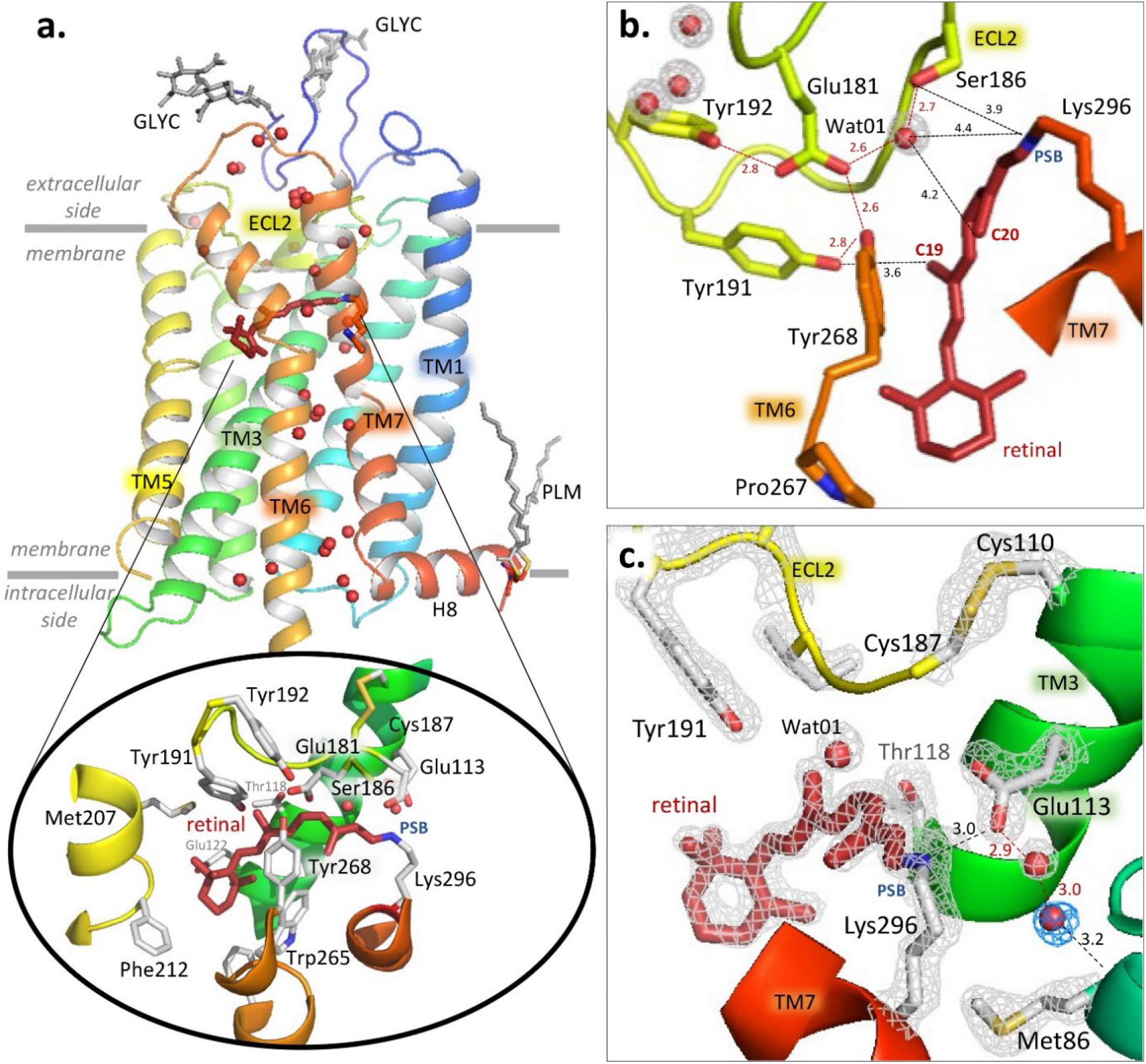
Room-temperature serial femtosecond crystallographic (SFX) structure of the dark state of bovine rhodopsin from crystals grown in lipidic cubic phase. **a.)** Overall structure of rhodopsin rainbow-colored by residue number from blue (N-terminus) to red (C-terminus). The seven-transmembrane bundle contains two N-glycosylation domains (GLYC) and palmitate groups (PLM) that anchor the amphipathic helix H8 to the membrane (grey lines). Water molecules (red spheres) form key networks^2^ between the extracellular (retinal ligand binding pocket) and intracellular (G protein binding site) regions of the receptor. The 11-cis retinal (dark red) is covalently bound to Lys296 (see inset) via a protonated Schiff base (PSB). The retinal binding pocket is further composed of amino acids surrounding respectively the PSB (the counterion Glu113 and Met44, Phe91, Thr94, Ala292, Phe293), the retinal aliphatic chain (Ala117, Thr118, Tyr191, Trp265 and Glu181/Ser186 through water Wat01) and the β-ionone ring (Glu122, Phe212, Met207, Phe261, Ala269; for clarity, only selected residues in the binding pocket are shown). **b-c)** Examples of well-resolved molecules in the water-mediated networks connecting **(b)** the counterion Glu113 to Met86 of TM2 (and Ala117^<u>3.32</u>^, not shown) and **(c)** the residues in the ancestral counterion Glu181 network. The water molecules have well-defined electron densities (grey and blue meshes, 2Fobs-Fcalc electron density contoured at 2.2 and 0.7 σ, respectively). Presence of the newly resolved water WAT04 contoured in blue.

In recent years, time-resolved crystallography^16,17^ at X-ray free-electron lasers (XFEL) has been used to reveal ultrafast structural changes in myoglobin (as carbon monoxide photo-dissociates from the haem cofactor^18^), photoactive yellow protein^19^ and bacterial phytochromes^20^ (in response to the isomerisation of the co-factors *p*-coumaric acid and biliverdin respectively), microbial proton^21,22,23^, sodium^24^ and chloride^25^ pumps (bacteriorhodopsin, KR2, and NmHR respectively, in response to retinal isomerisation), and a bacterial photosynthetic reaction centre^26^. The protein molecules in the crystals are photoactivated with an optical laser pulse and the structure probed with an X-ray pulse from an XFEL, after a certain time delay. Each crystal generates one diffraction pattern, from a certain, random orientation. Hence, the experiment is carried out in a serial way: microcrystals are brought into the interaction region by using a continuous jet, and frames are collected from tens to hundreds of thousands of randomly oriented crystals.

In this work, we use time-resolved serial femtosecond crystallography (TR-SFX) at room temperature to follow the light-induced conversion of the inverse agonist 11-cis retinal to an agonist all-trans conformation in a vertebrate opsin. Our observations reveal how this translates into early structural changes within the protein. After 1 picosecond, we observe a twisted retinal that stores energy while structural motions in the protein radiate as anisotropic propagation away from the retinal chromophore. The rhodopsin structure, 100 ps later, reveals a slightly more relaxed conformation.

## A room temperature structure of rhodopsin obtained by serial femtosecond crystallography

The room-temperature SFX structure of rhodopsin in the inactive dark state was obtained to a resolution of 1.8 Å (**Extended Data Table 1, dark state**) from micro-crystals grown in a lipidic cubic phase (LCP). As with most membrane protein structures determined from LCP-grown crystals, these crystals display a Type I lattice^27^ forming 2D-layers of stacked protein built through hydrophobic interactions (**Extended Data Figure 1**). Potentially physiologically relevant dimers of rhodopsin molecules^28^ form contacts between the TM1 and helix 8 (H8) segments of each monomer and are assembled in a head-to-tail manner generating the asymmetric unit. By close inspection of the diffraction data and the resulting electron density maps, the presence of translation-related crystal domains was detected. Measured intensities were corrected to account for this, globally improving the quality and interpretability of the maps (**Methods**, **Extended Data Figure 2 and Table1)**. Overall, the SFX structure in the inactive dark state of rhodopsin (**Fig. 1a**) is very similar to other crystal structures collected at cryogenic temperatures (e.g., 1GZM^5^; RMSD = 0.33 Å on Cα atoms). In contrast with earlier structures solved in cryogenic conditions, the present room-temperature structure reveals electron density for all the previously described functional and structural water molecules. These include the water-mediated cluster around the ancestral counterion Glu181^29^ and its polar tyrosine cage (**Fig. 1b**), which play a role later in the photoactivation process^30,31^. Moreover, a new ordered water molecule was resolved near the Schiff base connecting the proximal counterion Glu113^<u>3.28</u>^ to Met86^<u>2.53</u>^ and Ala117^<u>3.32</u>^. This interaction has a central position at the TM2-TM3-TM7 interface in the transmembrane bundle (**Fig. 1c**).

## A bent structure of retinal after 1 picosecond of rhodopsin photoactivation

The first metastable intermediate of rhodopsin (bathorhodopsin, Batho-Rh)^15,33,34^ arises 200 fs after photo-activation. It is fully populated by Δ*t* = 1 ps^14,35^ and persists for tens of nanoseconds^36,37^. To characterize the structure of Batho-Rh we collected TR-SFX data at the Swiss and Japanese XFELs (**Extended Data Figure 3)** from LCP-grown microcrystals of rhodopsin photo-activated using a femtosecond-pump laser with a 480 nm wavelength for three time-delays of photoactivation of Δ*t* = 1 ps, 10 ps and 100 ps. High-quality TR-SFX data (**Extended Data Table 1**) are represented as difference Fourier electron density maps in **Extended Data Figure 5**. For the shorter time-delay, changes in electron density are highly anisotropic, clustering in the immediate vicinity of the buried retinal chromophore and propagating towards the cytoplasmic side of the protein via the TM5 and TM6 helices at a minimum speed of 18Å/ps measured along the TM6 (1800 m/s, slightly above the speed of sound in water). This structural anisotropy had completely decayed by Δt = 100 ps (**Extended Data Figure 5)**.

Light-induced structural changes within the retinal polyene chain are observed at Δ*t* = 1ps as a strong negative difference electron density feature (minimum of −6.2 σ, where σ is the root mean square electron density of the unit cell) and a complementary positive difference electron density feature (maximum of +5.8 σ) **(Fig. 2a)** associated with the C11=C12 double bond revealing that this bond has isomerized. This event is associated with changes in electron density near the C20 methyl (−6.6 σ and +5.6 σ). Modelling these electron density changes in combination with structural refinement against crystallographic observations extrapolated to 100% occupancy of the photo-activated intermediate (for photoactivation levels, see **Methods**)(**Extended Data Table 1; 1, 10 and 100 ps time delays**) establish that retinal isomerisation involves a large clockwise rotation (viewed from the PSB) of the C20-methyl (47.7°) towards the extracellular side in concert with a shift of both the proximal water molecule WAT01 and Tyr268^<u>6.51</u>^ of the Glu181 cage **(Fig. 2b)**. This tilt of the C20-methyl increases to 51.3° by Δ*t* = 100 ps (**Extended data 6**). The corresponding distortion was measured as 54.9° in a cryo-trapped bathorhodopsin intermediate study that lacked any temporal resolution^10^. The plane of retinal containing the C19-methyl, which is located on the opposite side of the isomerising C11-C12 bond, is fixed in the resting state by Thr118^<u>3.33</u>^ (3.36 Å away)^38,39^ and Tyr191^<u>ECL2</u>^. Consequently, the C19 methyl is only minimally affected by the cis-to-trans isomerisation (**Extended data 6**), shifting of only half an angstrom towards Tyr191 (36.0° rotation compensated by a backward elbow movement of the polyene chain) and in the opposite direction to the C20 methyl **(Fig. 2b-c)**.

**Fig. 2.**
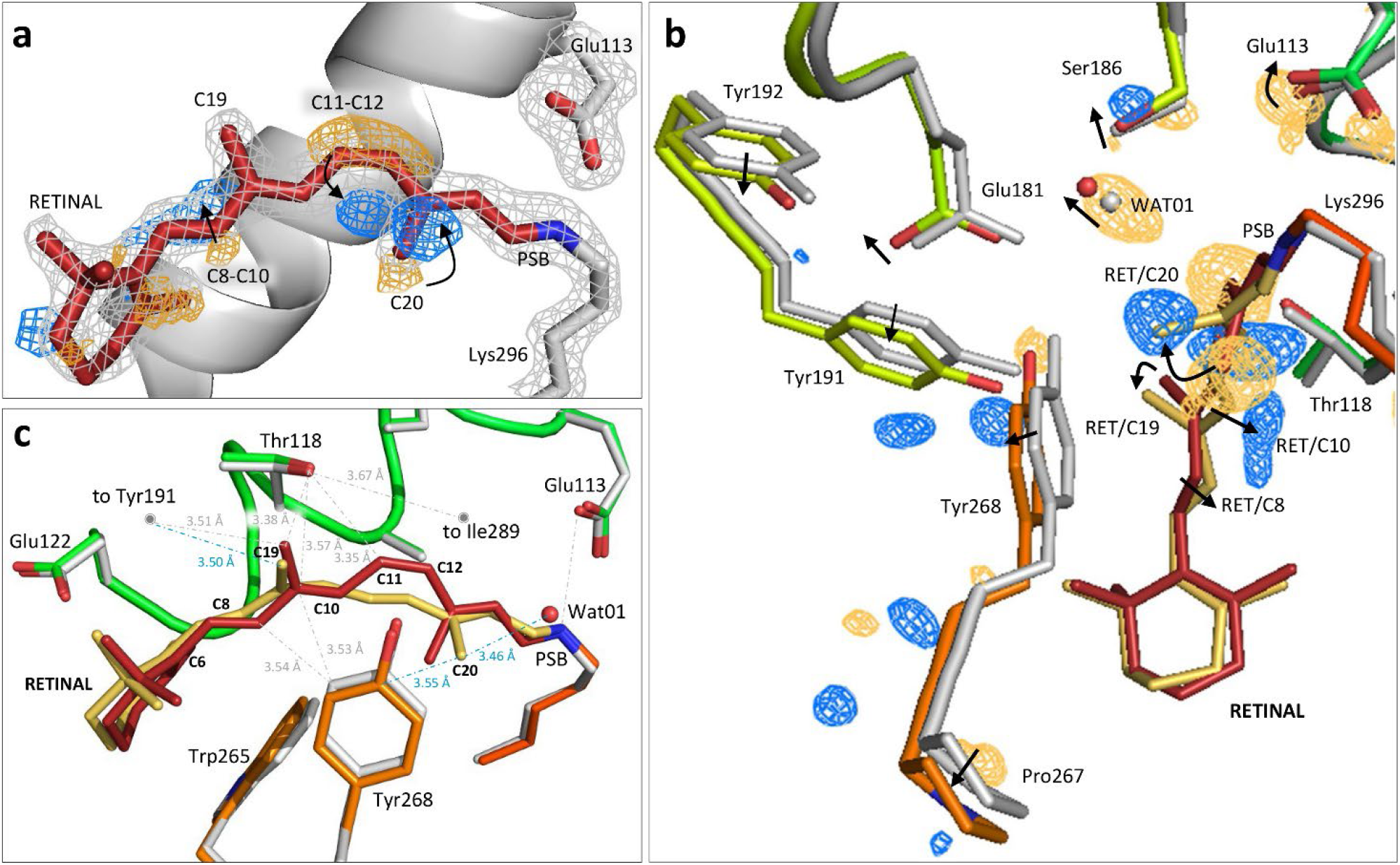
Retinal conformation captured 1 picosecond after rhodopsin photoactivation using TR-SFX. **a)** Depiction of the electron density changes. The retinal model (in red) and the contoured grey mesh (at 2.7σ of the 2F_obs_-F_calc_ electron density map) correspond to rhodopsin in the dark state obtained by SFX. The difference Fourier electron density (Fobs^light^ - Fobs^dark^ contoured at 3.8 σ) around the C11=C12 bond of the polyene chain and the C20-methyl show features appearing after 1 ps photoactivation in blue (positive density) that are correlated with disappearing features in gold (negative density), establishing that the chromophore has already isomerised. A positive density is also observed along C8 and C10 of the retinal polyene chain. **b-c)** Effect of retinal isomerisation on the surrounding amino acid residues. The model of 1 ps-photoactivated rhodopsin (retinal in yellow; rhodopsin in orange and green) obtained from the extrapolated map 2Fext - Fcalc (21% photoactivation, see **Methods**) superimposed to the dark state model (retinal in red; rhodopsin in grey). The main chain Cɑ atoms of the protein were used for the structural superposition. The difference electron density map **(b)** (Fobs^light^ - Fobs^dark^, contoured at 3.4 σ) shows the presence of positive and negative electron densities (blue and yellow) around specific amino acids such as Tyr268 of the binding pocket. Arrows illustrate shifts or rotations. Panel **(c)** illustrates the torsion of the retinal polyene chain at C11-C13 in the direction of Tyr268 and the bending along C6-C11. Selected distances from retinal to rhodopsin residues are depicted as grey dotted lines for the dark state and as blue dotted lines for the isomerised form.

These rearrangements in the retinal molecule at Δ*t* = 1 ps are compatible with an aborted ‘bicycle pedal’ mechanism of photoisomerisation^14,40^ in which the interactions of the C19 methyl of retinal with specific residues in the tight binding pocket confer resistance to a larger rotation of this methyl group. The consequence for such aborted C9=C10 isomerisation is the release of energy over the polyene chain, affecting mostly C8 and C10 which elbow in the opposite directions relative to the C20 methyl **(Fig. 2b-c)**. The C6-C11 segment of the retinal polyene chain is bent, aligning all carbons in a nearly perfect arc **(Fig. 2c)** affecting the surrounding interactions in the binding pocket.

The absorption maximum of this state was computed by quantum mechanics/molecular mechanics (QM/MM) optimization of the experimental structure at Δ*t* = 1 ps and a subsequent vertical excitation energy calculation between the ground and the optically active first excited state of the PSB (see **Methods**). This yielded a value of 522 nm, which is in good agreement with the experimental absorption maximum of 529 nm^41^ for Batho-Rh (**Extended Data Table 2**). Our QM calculations also show that the twisted all-trans retinal at 1 ps stores an excess of 20 kcal/mol of energy (for comparison, the energy of a 480nm photon is 59.6 kcal/mol) compared to the planar 11-cis conformation, which is in qualitative agreement with the experimental value of 32 kcal/mol measured for the rhodopsin-to-Batho-Rh transition^42^.

## Picosecond changes in the rhodopsin binding pocket

Light-induced isomerisation transforms retinal into an agonist that interacts differently with residues in the rhodopsin binding site. One picosecond after photoactivation, the isomerised all-trans retinal fills the same volume as the 11-cis resting conformation but is now free from several hydrogen bonds and van der Waals interactions that stabilize the dark state structure in an inactive conformation **(Fig. 3a)**. The covalently bound retinal becomes bent like an arc that is stabilized in the middle by interactions between the C19- and C20-methyl groups of the retinal and two tyrosine residues from the ancestral counterion network (Glu181 polar cage), Tyr191^<u>ECL2</u>^ and Tyr268^<u>6.51</u>^. The isomerisation and rotation of the C20-methyl-C13-C14 plane induce a kink of the C15 from the polyene chain influencing minimally the neighbouring PSB/Glu113^<u>3.28</u>^ salt bridge, but the surrounding water WAT04 order of the Glu113 hydrogen bond network **(Fig. 2c and Extended Data Figure 7)**. The interaction between the polyene chain and Ala117^<u>3.32</u>^-Thr118^<u>3.33</u>^ are two critical retinal contacts with TM3 that are weakened at Δ*t* = 1 ps **(Fig. 3b-c, Extended Data Figure 8)**. An additional interaction with TM3 is disrupted, between the isomerising bond and Cys187^<u>ECL2</u>^ of the structurally important and highly conserved disulphide bridge Cys187-Cys110, linking TM3 to ECL2^43^ **(Fig. 3a-b)**. Remarkably, the light-induced structural changes observed around the retinal of bovine rhodopsin bear some resemblance to those observed in other low-homology seven-transmembrane helix retinal-binding proteins from bacteria and archaea. For instance, a weakening or disruption of the interaction between retinal and TM3 (which corresponds to Helix C in prokaryotic opsins) is also observed in bacteriorhodopsin (bR)^23^, the sodium photosensitive pump KR2^23,24^, and the chloride pump *Nm*HR^44^, which all undergo completely different activation mechanisms. Thus, despite their different evolutionary origins and types of isomerisation (11-cis-to-all-trans versus all-trans-to-13-cis), retinal-binding proteins seem to share the need to disengage the retinal from the central transmembrane helix before undergoing the next steps of activation.

**Fig. 3.**
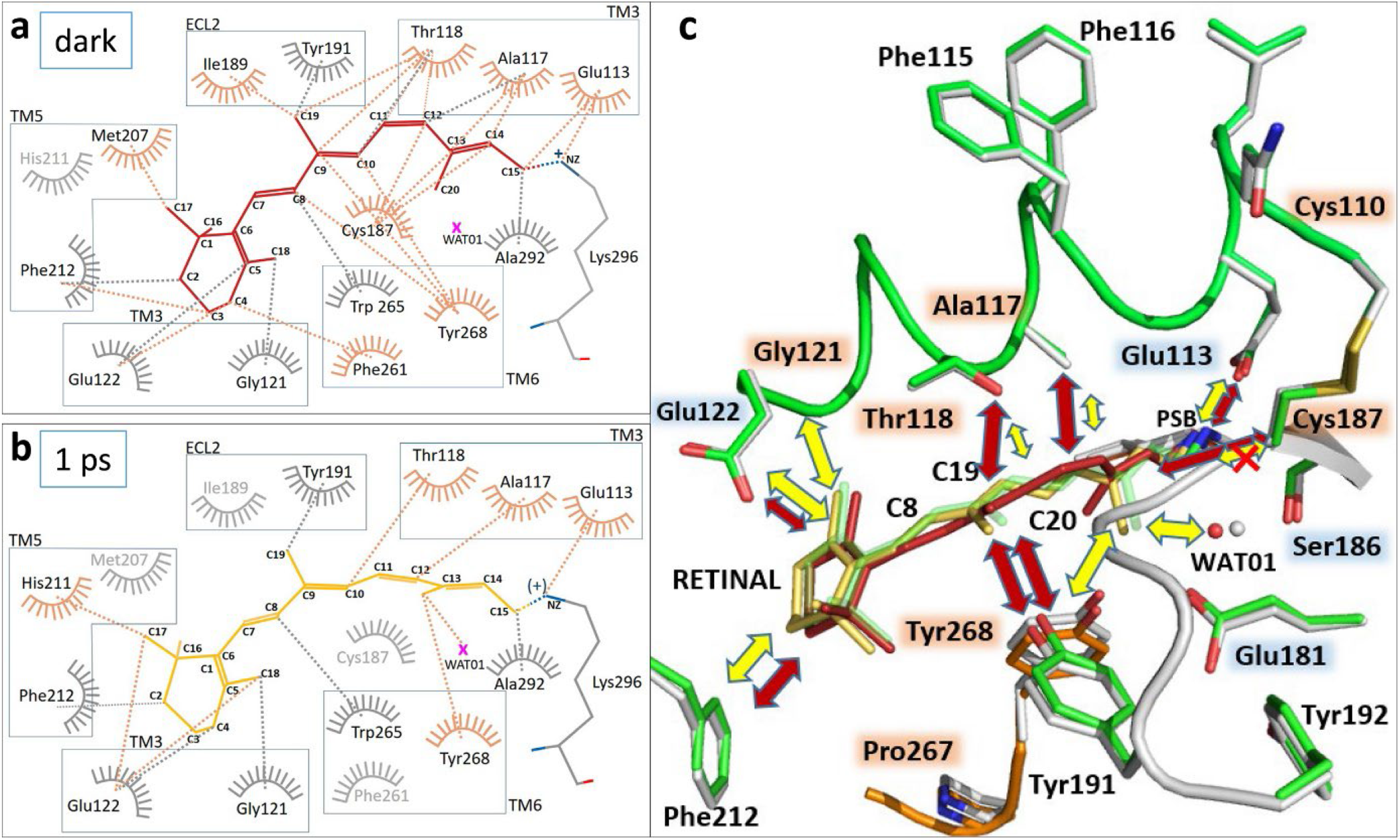
Interactions of retinal with the binding pocket residues are dramatically reduced 1 picosecond after photoactivation. **a-b.)** LIGPLOT representations of the interactions of 11-cis retinal with the binding pocket in the dark state (**a**) and of all-trans retinal after 1 ps of photoactivation (**b**). For a comparison with the 10 and 100 ps structure, see **Extended Data Figures 6 and 8**. All interactions closer than 3.7 Å are displayed. Grey color is used for interactions that do not change. Orange color highlights amino acids and interactions of interest. The pink cross represents WAT01. **c.)** Schematic representation of the interactions of retinal in the rhodopsin ligand binding pocket before (red arrows) and after photoactivation in the picosecond range (yellow arrows). A larger arrow means stronger interaction. The grey structure corresponds to the dark state (retinal in red) and the colored structure corresponds to the 1ps illuminated model (retinal in yellow). For comparison, the retinal model after 100ps is shown in translucent green.

As mammalian rhodopsin evolves along its reaction pathway, some of the observed changes will become more pronounced while others will revert (**Extended Data Table 3**). By Δ*t* = 10 and 100 ps, the β-ionone ring and C19-C20 methyl groups still interact respectively with Gly121^<u>3.36</u>^-Glu122^<u>3.37</u>^ and the tyrosines of the Glu181 polar cage **(Extended Data Figure 8)** while e.g. Tyr268 and Pro267 relax to their initial positions.

Whereas rhodopsin is considered a prototypical Class A GPCR, the mechanism used by GPCRs to recognize diffusible agonist ligands by conformational selection is vastly different from the extreme case of induced-fit displayed by light-activated GPCRs such as rhodopsin^45^. Remarkably, both conformational selection and induced fit converge rapidely into a common GPCR activation mechanism^46^ and thus the early stages of retinal isomerisation may reveal fundamental determinants of agonism in GPCRs. For instance, early structural changes in the retinal binding pocket are associated with a small outward tilt (~0.5 Å) near Pro215^<u>5.50</u>^ and Pro267^<u>6.50</u>^, which are located in the middle of TM5 and TM6 **(Fig. 2b)** and enable anisotropic motions in the extracellular part of the receptor (**Extended Data Figure 5**). Both prolines are conserved in Class A GPCRs and are key in agonist-induced activation^47^. Moreover, TM3 is a region that plays a central role in the architecture of the transmembrane bundle of Class A GPCRs^48^ by forming part of both the ligand and the G protein binding pockets. Our TR-SFX data reveal that, even at an early stage of activation, the inverse agonist (11-cis retinal) has stripped itself away from TM3 while isomerising into the agonist conformation **(Fig. 3b)**. Several of the affected positions in TM3 (Glu113^<u>3.28</u>^, Thr118^<u>3.33</u>^, Gly121^<u>3.36</u>^, and Glu122^<u>3.37</u>^) correspond to conserved residues in the binding site of Class A GPCRs that are involved in ligand binding^48^, with Gly121^<u>3.36</u>^ in particular being part of a consensus ‘cradle’ scaffold for ligand recognition^48^. Our TR-SFX observations using XFEL thus reveal how the photoactivated conformation of the retinal rapidly weakens many weak interactions with the amino acids of the rhodopsin binding pocket **(Fig. 3c)** and thereby commits the receptor’s relaxation pathway towards its G-protein binding signalling conformation.

## Conclusions

Our high-resolution SFX structure of rhodopsin in the inactive dark state at room-temperature reveals the entirety of the water-mediated hydrogen bond network within the protein. One picosecond after light activation, rhodopsin has reached the red-shifted Batho-Rh intermediate. Already by this early stage of activation, the twisted retinal is freed from many of its interactions with the binding pocket while structural perturbations radiate away as a transient anisotropic breathing motion that is almost entirely decayed by 100 ps. Other subtle and transient structural rearrangements within the protein arise in important regions for GPCR activation and bear similarities to those observed by TR-SFX during photoactivation of seven-transmembrane helix retinal-binding proteins from bacteria and archaea. We thus suggest that the protein disperses an initial excess of energy through the early GPCR structural pathways that will be used for activation. Our study reveals an ultrafast energy dissipation in rhodopsin occuring via conserved residues of GPCR activation pathways and lays the experimental groundwork to study the early activation events in the large family of Class A GPCRs.

## Acknowledgements

This project has received funding from the European Union’s Horizon 2020 research and innovation programme under the Marie Skłodowska-Curie grant agreements No 701647 (to M.R.), 701646 (to S.Br.) and 884104 (PSI-FELLOW-III-3i) to S.S. Also from the Swiss National Science Foundation through an “Ambizione” grant PZ00P3_174169 (to P.N) and project grants 31003A_179351 (to J.S.) and 192780 (to X.D.), and 310030B_173335 (to G.F.X.S.), and through the NCCR:MUST program (to C.M. and J.S.). F.P. acknowledges ETH Zürich through the National Center of Competence in Research Molecular Ultrafast Science and Technology and the ETH Femtosecond and Attosecond Science and Technology programs. The project is supported by the Japan Society for the Promotion of Science KAKENHI Grant No.19H05776 # supported by Platform Project for Supporting Drug Discovery and Life Science Research (Basis for Supporting Innovative Drug Discovery and Life Science Research (BINDS)) from AMED under Grant Number JP21am0101070 to S.I. This work was supported by the National Science Centre (Poland) under grant No. 2017/27/B/ST2/01890 to A.W. RN acknowledges financial support from the Swedish Research Council (grant No. 2015-00560)

We thank members of the Engineering Team of RIKEN SPring-8 Center for technical support. XFEL experiments were conducted at BL3 of SACLA with the approval of the Japan Synchrotron Radiation Research Institute (Proposal Numbers 2015B8043 and 2018A8066). We acknowledge the SwissFEL and the Swiss Light Source synchrotron for excellent performance and support from the Alvra team and Macromolecular Crystallography group, respectively, during the beamtimes allocated for this study. The TR-WAXS experiment was performed at the Linac Coherent Light Source (LCLS), SLAC National Accelerator Laboratory. Use of the LCLS is supported by the U. S. Department of Energy, Office of Science, Office of Basic Energy Sciences under Contract No. DE-AC02-76SF00515. Part of the sample injector used at LCLS for this research was funded by the National Institutes of Health, P41GM103393, formerly P41RR001209.

We thank Tomas White (CFEL, Hamburg) and Takanori Nakane for the advices regarding the use of the CrystFEL software and the SACLA processing pipeline, respectively. We thank Timm Maier (University Basel) for access to the SONICC imager used for finding the first rhodopsin crystal hits. We thank the members of the High Performance Computing and Emerging Technologies Group at the Paul Scherrer Institute for technical support with the QM/MM simulations. We thank Cornelius Gati for initially helping to measure some rhodopsin crystals controls at the SONICC instrument (Hamburg). We thank Gregor Cicchetti for his comments on the paper.

## Author contributions

The project was initiated by G.F.X.S. with input from R.N. and coordinated by V.P. G.F.X.S. coordinated and supported crystallographic applications at SwissFEL and contributed to discussions throughout the project. So Iwata, Eriko Nango and Rie Tanaka shared experience, coordinated and supported crystallographic applications at SACLA. Initial XFELs pilot tests at the LCLS and SACLA were supported by S.Bo. and K.T.+Y.J.+S.O. respectively, in a beamtime of J.S. S.Bo. prepared the X-ray instrument and collected the data. Initial rhodopsin crystals were developed in an experiment with T.G., P.M. and V.P. Highly diffracting crystals were optimized and tested for diffraction at the Swiss Light Source by T.G., P.S. and V.P. ROS membranes from bovine retinae were purified by C.J.T., W.W., V.P. and T.G. Rhodopsin was purified and crystallized by T.G., with supplementary support before beamtimes by A.F., S.M., C.J.T., P.M., J.M., F.P., N.V., W.W. and V.P. Pump–probe experiments at the Alvra endstation of the SwissFEL were prepared by C.M., P.J., C.Ci and D.J. J.S., R.N. and D.J. supported V.P. in setting the laser power and scheme of data collection. The lipidic sample injection was optimized by T.G., N.V. and D.J. Sample preparation and reservoir loading in darkness at the XFEL was done by T.G., A.F., S.M., G.O., D.K., E.L, H.G., B.P., I.M. and injector for viscous samples were operated during the XFEL beamtimes by G.G., P.B, D.G., O.T., A.W. and D.J., F.D., respectively. The endstation including the laser system was aligned and operated by P.J., G.K., C.Ci., C.B. and C.M., who also designed the Alvra prime pump-probe station. The SFX data analysis pipeline of SwissFEL was built and operated by D.O. and K.N. Data processing during the beamtime was done by T.W., P.N., K.N. with the help of D.O., B.P., D.K., S.Br. and C.Ca. D.K., X.D., C.Ca, A.F., V.P. recorded progress during data collection. Data processing was further completed by T.W., T.G., K.N., M.R., V.K., A.D., C.Ca. Crystal lattice translation was identified and corrected by M.R., C.Ca., V.P. and T.W. Structures were refined by T.G., M.R., T.W. and data interpreted by V.P with the help of X.D. and G.F.X.S. Quantum mechanics/molecular mechanics calculations and Quantum-chemical calculations were done by S.S. and X.D. The TR-XSS experiment was designed by R.N., G.F.X.S. and analysed by D.S., R.N. The manuscript was written by V.P. with direct contributions from T.G., X.D., G.F.X.S and R.N. and with further suggestions from most of the other authors. All authors read and acknowledged the manuscript.

## Data availability

Coordinates and structure factors have been deposited in the Protein Data Bank under accession codes 7ZBC (SFX dark state rhodopsin at the SACLA), 7ZBE (SFX dark state rhodopsin at the SACLA) and 8A6C, 8A6D, 8A6E for the photoactivated rhodopsin with time-delays 1ps, 10ps and 100ps, respectively.

## METHODS

### Rhodopsin extraction from retinae and purification

All extraction and purification steps were carried out under dim red light conditions. Commercially available dark-adapted frozen bovine retinae (W. L: Lawson Company, USA) were used to isolate rod outer segment (ROS) membranes following the protocol described in Okada et al. 1998^49^. In short, 200 frozen bovine retina were diluted in ROS buffer (10 mM MOPS, 30 mM NaCl, 60 mM KCl, 2 mM MgCl_2_, 1 mM DTT), 40 % (w/w) sucrose and two tablets of cOmplete™ Protease Inhibitor Cocktail. The mixture was shaken by hand for 4 min, centrifuged at 4 °C, 4’000 g for 30 min. This step was repeated and the pooled supernatants were combined, diluted by a half with ROS buffer containing no sucrose and centrifuged at 4 °C, 24’000 g for 30 minutes. Pellets were resuspended in ROS buffer containing 23.4 % sucrose and layered on a freshly prepared gradient with two layers of ROS buffer with 34 % (w/w) and 29 % (w/w) sucrose. The ROS membrane-loaded gradients were centrifuged using a swing-out rotor SW28 at 4 °C at 110’000 g for 90 minutes and rhodopsin containing layers (23-29 % interface and 29 % layer) were aspirated and flash frozen in liquid nitrogen. The rhodopsin concentration of the dark state ROS membranes, determined by recording a UV/VIS spectrum before and after illumination, yielded commonly 200 ± 40 mg rhodopsin. Bovine rhodopsin can be further isolated by detergent solubilisation and affinity chromatography using a Concanavalin A resin (ConA, GE Healthcare Life Sciences, USA) as described in Edwards et al. 2004^50^. This protocol was optimized: (I) the starting retinae material was doubled; (II) the amount of resin was scaled-up to trice; (III) in order to sharpen the elution profile the second half of the elution phase was performed in reversed flow. A ROS membrane suspension containing around 180±20 mg rhodopsin was diluted three times in ConA buffer (50 mM NaOAc, 150 mM NaCl, 3 mM MgCl_2_ 6H_2_O, 3 mM MnCl_2_ 4H_2_O, 3 mM CaCl_2_ 2H_2_O, 1 mM Na2-EDTA 2H_2_O, 2 mM 2-mercaptoethanol, pH 6) and spun down at 4 °C, 104’000 g for 35 minutes. The resulting ROS membranes pellet was resuspended in 90 mL ConA buffer containing one tablet of protease inhibitor and the membranes were solubilized at room temperature with lauryldimethylamine-oxide (LDAO, Sigma-Aldrich, USA) to a final concentration of 1 %. The solubilized sample was spun down at 4 °C, 118’000 g for 60 min before Concanavalin A affinity chromatography in ConA buffer containing 0.1 % (w/v) LDAO and direct elution with 0.2 M methyl α-D-mannopyranoside in the same buffer. Aliquots of rhodopsin at 2 mg/mL were flash frozen in liquid nitrogen and ready to use for crystallization in lipidic cubic phase.

### Crystallization and TR-SFX sample preparation

All crystallization and sample preparation steps were carried out under dim red light conditions. Prior to crystallization in lipidic cubic phase (LCP) flash-frozen aliquots of purified rhodopsin were thawed and the detergent was exchanged for 0.21% n-Decyl-N,N-Dimethylamine-N-Oxide (DAO, Anatrace, Inc., USA) in 50 mM Na Acetate, 150 mM NaCl, 3 mM MgCl2, pH 6.0 using a PD10 buffer exchange column (Sigma-Aldrich, USA). The eluate was spun down at 18 °C, 21’000 g for 15 min and further concentrated to at least 20-25 mg/ml using a concentrator Ultra 4 MWCO 30 at 18 °C, 4’000 g. After centrifugation at 18 °C, 21’000 g for 15 min, the protein sample was mixed at 22 °C with monoolein (1-oleoyl-rac-glycerol, Nu-Check prep) in a ratio of 40 to 60, respectively, using gas-tight Hamilton syringes until formation of the translucid LCP. The well diffracting rhodopsin crystal was initially obtained from a low molecular weight polyethylene glycol screen with various types of buffer and the first hit detected using the second harmonic generation imaging SONICC^®^ (second order nonlinear optical imaging of chiral crystals) device (Formulatrix)^51^. For crystal growth, 20-40 μl of protein-laden LCP were injected into precipitant laded Hamilton glass syringes containing 200-400 μl of precipitant (37-39 % PEG 600 and 100 mM Bicine pH 9.0). Alternatively, 80 μl of protein-laden LCP were injected into 1 ml plastic syringes containing 800 μl of precipitant. The samples were wrapped in aluminium and stored at 18 °C. After 3 days, plate-shaped crystals in LCP with dimensions of about 15 x 15 x 1.5 μm can be harvested by removing the precipitant and kept stable for weeks in the darkness at 18 °C. At the X-ray free-electron laser beamline, the LCP sample containing rhodopsin crystals was imperatively mixed with a 3 way-coupler to ensure homogeneity ^52^ prior to loading into a reservoir of the high viscosity injector. In case of residual precipitant contamination, the LCP-laden crystal sample was titrated with PEG 1,000 (50 % (w/v)) and finally mixed 1 : 5 with monoolein.

### Time-resolved pump probe serial crystallography

X-ray diffraction data were collected at the X-ray free-electron lasers (XFELs) BL3_EH2 end station of the SACLA^53^ (beamtimes 2015B8043 and 2018A8066) and Alvra end station of the SwissFEL (beamtimes 20172060 and 20200597). The energy of the X-ray beam was 9-10 keV with a pulse length of 10 fs (SACLA) and 65 fs (SwissFEL) and a focus at the sample of 1 × 1 μm (SACLA) and 5 × 5 μm (SwissFEL). The hutch was prepared for dim red light conditions and the femtosecond pump laser set at a wavelength of 480nm with a pulse energy of 9 (SACLA) to 5 (SwissFEL) μJ/100 fs pulse duration at the sample position. The pump laser beam size was of 47-50 μm FWHM (80-85 μm 1/e^2^). If one assumes idealized Gaussian beam optics then this corresponds to a peak energy density of 200 mJ/cm^2^ at SwissFEL (Δ*t* = 1 ps) or alternatively a mean energy density of 140 mJ/cm^2^ averaged over the laser’s focal FWHM (compare with tabulated data in reference^17^; or a peak power density of 2000 GW/cm^2^. The corresponding values for the SACLA study (Δ*t* = 100 ps) are a peak fluence of 360 mJ/cm^2^, and average fluence of 260 mJ/cm^2^, and a peak power density of 3600 GW/cm^2^. If one calculates the product of the average fluence (F) with the resting state absorption cross-section (σ) divided by the energy of a single photon (h‧ν, where h is Planck’s constant) we recover σ‧F/h‧ν = 45 for the illumination conditions at SwissFEL and σ‧F/h‧ν = 81 for those used at SACLA. Whereas these values may suggest considerable multiphoton excitation, the results from time-resolved X-ray solution scattering studies on rhodopsin (**Extended data Figure 4**) imply that far fewer photons are absorbed (see Time-resolved X-ray solution scattering below).

Crystals of bovine rhodopsin grown in LCP were used to collect time-resolved serial femtosecond crystallography data at the time delays of 1 ps, 10 ps (SwissFEL) and 100 ps (SACLA)(**Extended Data Figure 6)**. The crystals where extruded using a high viscosity injector through a 75 μm nozzle with a constant flow rate of 0.033 μl/min (SwissFEL) or 2.5 μl/min (SACLA)^54^ to the pump probe intersection point where the data were collected with every fifth shot of the pump laser blocked (data collection scheme of 4 light-activated, then 1 dark)(SwissFEL) or interleaving ON/OFF-laser (collection in the mode 1 light: 1 dark (SACLA)), depending on the repetition rates of the XFELs (SwissFEL 25 Hz; SACLA 30 Hz) and pump lasers (SwissFEL 25 Hz; SACLA 15 Hz), respectively. As a control, a true dark state data has been also collected with the pump laser off (all dark data, SFX mode).

### Time-resolved X-ray solution scattering

Time-resolved X-ray solution scattering (TR-XSS) studies using samples of detergent solubilized rhodopsin and a liquid microjet for sample injection were performed at the LCLS as previously described^55^. Rhodopsin was solubilized in DDM (n-Dodecyl-β-Maltoside) to a concentration of 8.4 mg/ml (0.2 mM). Samples were photoactivated using 480 nm laser pulses 50 fs in duration, focused through a 1/e^2^ spot 100 μm in diameter (59 μm FWHM) with pulse powers of 6 μJ (110 mJ/cm^2^ averaged across the FWHM; 3000 GW/cm^2^ peak power), 22 μJ (400 mJ/cm^2^ averaged across the FWHM; 11200 GW/cm^2^ peak power), 45 μJ (830 mJ/cm^2^ averaged across the FWHM; 22900 GW/cm^2^ peak power) and 89 μJ (1640 mJ/cm^2^ averaged across the FWHM; 45300 GW/cm^2^ peak power) respectively. Laser induced heating was estimated from these data from the time-delays 10 ps ≤ Δt ≤ 1 μs (**Extended Data Figure 4a**) since TR-XSS studies on detergent solubilized photosynthetic reaction centre have shown that there is relatively little change in the heating signal over this time-domain^55^. The principal singular value decomposition (SVD) component from these data was compared with temperature calibration curves (**Extended Data Figure 4b**) in order to estimate the laser-induced change in temperature for different photoexcitation fluence, as described in reference^55^. The heating impulse thus imparted is shown in **Extended Data Figure 4c**, with a negative time-point used as a control (plotted as zero laser fluence).

Using **Extended Data Figure 4c** as a laser heating-induced calibration curve, it was estimated that the temperature jump induced under the photo-excitation conditions at SwissFEL (fluence of 140 mJ/cm^2^ averaged over the FWHM) would be ΔT = 0.016 °C ± 0.009 °C; and the photo-excitations used at SACLA (fluence of 260 mJ/cm^2^ averaged over the FWHM) would induce a temperature jump ΔT = 0.028 °C ± 0.016 °C. This corresponds to an excess 1.2 ± 0.7 photons absorbed by the rhodopsin chromophore under the conditions at SwissFEL, and 2.1 ± 1.2 photons under the conditions at SACLA. Although considerable uncertainly must be acknowledged in these estimates, these experimental values are strikingly lower than those calculated as σ‧F/h‧ν = 45 for the photoexcitation conditions used at SwissFEL and σ‧F/h‧ν = 81 for the photoexcitation conditions use at SACLA. There has been considerable debate about what constitutes appropriate photo-excitation conditions for TR-SFX studies^17, 56, 57, 58^. Our TR-XSS observations suggest that laser induced sample heating is much lower than expected and this is consistent with results from other TR-XSS studies on other light sensitive proteins^17^. This may be due to the cross-section of the first excited state at 480 nm being much lower than that of the ground state; may be due the excited states having relatively high stimulated Raman scattering and stimulated emission cross-sections and therefore the absorbed excess energy is carried away by emitted photons rather than ultimately emerging as sample heating; may be due to high scattering from the microjet causing most of the incoming light from the laser pulse to be lost; or may be due to a combination of all of these factors and others. Nevertheless, these observations strongly imply that it is unlikely that our TR-XSS data are dominated by the quasi-isotropic structural effects of laser induced heating, as observed for a photosynthetic reaction centre when the order of 800 photons were absorbed per chromophore^55^.

### Data processing

All data were indexed using INDEXAMAJIG with the XGANDALF algorithm for data collected at SwissFEL (SF dark, 1 ps, 10 ps) and the MOSFLM^59^, DirAx^60^ and XGANDALF^61^ algorithms for data collected at SACLA (SACLA dark, 100 ps). The integration radius was set to 2 pixels for SwissFEL data and 3 pixels for SACLA data while the background annulus was set to between 4 and 6 pixels for SwissFEL data and 4 and 7 pixels for data collected at SACLA. The crystal to detector distance was optimised on a per-run basis by sampling detector distances between 91.5 mm – 97.5 mm (SwissFEL) and 47.5 mm – 53.5 mm (SACLA) first at 200 μm and then 20 μm increments to determine the detector distance at which the standard deviations of the unit cell dimensions were minimised.

The SwissFEL and SACLA data were scaled and merged separately in PARTIALATOR^62^, using partiality modelling with XSPHERE^63^. Custom splitting was used to output separate dark and light activated reflection files for data collected at each free electron laser.

A lattice translocation defect was identified in the crystals after inspection of the Patterson map with phenix.xtriage^64^. In particular, the Patterson peak at td = (0.000, 0.245, 0.000) was attributed to the presence of two translation related domains in the crystals. The correction required to retrieve single-domain intensities is described in Wang et al., (2005)^65^, the percentage of molecules in the translated domain (κ) was determined by correcting the intensities at increasing κ values from 0% to 50% in 1% increments until the (0.000, 0.245, 0.000) Patterson peak was flattened. The correction led to a reduction in the SACLA dark state Rfree from 26.11% to 23.92% and improved the interpretability of the dark state electron density maps, which allowed further improvement of the model (**Extended Data Figure 2**).

### Structure determination and refinement of rhodopsin dark state

PDB entry with accession code 1U19^4^ with solvent and ligand molecules removed was used as a molecular replacement search model in *Phaser MR*^66^. The dark state structure was obtained after several interactive cycles of refinement and interactive model building with *Phenix.refine*^67^ and *Coot*^68^. An additional ligand geometry file was generated using JLigand to restrain the geometry of the protonated Schiff base linking the lysine side chain to retinal^69^.

### Calculation of difference density maps

Fo_light_ and Fo_dark_ amplitudes were calculated from the lattice translation defect corrected intensities using *phenix.french_wilson* ^64^ and Fo_light_-Fo_dark_ difference maps were calculated using *phenix.fobs_minus_fobs_map* ^64^ using the multi-scaling option excluding amplitudes smaller than 1σ and using reflections within the resolution range between 9 Å and 1.8 Å. All F_obs_(light)-F_obs_(dark) were computed using phases of the refined dark state.

### Data extrapolation

Extrapolated data were calculated using the lattice translation corrected data and following the method described by Pandey *et al*^70^. A linear approximation was used as follows: *F_extra_* = 100/*A* × (*F_obs_*(light) -*F_obs_* (dark)) + *F_calc_*, where *A* is the activation level in percent, *F_extra_* represents the extrapolated structure factor amplitudes and *F_calc_* represents the calculated amplitudes of the dark state model. The activation level for each time point was determined independently using the method described by Pandey et al^70^. Briefly, extrapolated data were calculated with activation levels ranging from 10% to 50% in 1% increments, and calculated 2*F_extra_* -*F_calc_* difference maps together with phases from the dark state model. Negative 2*F_extra_* - *F_calc_* density around C11, C12 and C20 of retinal, which display negative density features in the F_obs_ (light) - F_obs_ (dark) maps, was integrated with a radius of 1.5 Å and above at 1.5 σ cut-off for each activation level. Negative 2*F_extra_* - *F_calc_* difference density was plotted as a function of activation level, when the activation level is overestimated there is little 2*F_extra_* - *F_calc_* negative difference density at these atomic positions while the magnitude of the negative 2*F_extra_* - *F_calc_* difference density increases when the activation level is underestimated. The activation level is then determined by calculating the intersection between the two linear sections of the plot to find the activation level at which the negative density begins to appear. As the three light activated datasets were collected under different experimental conditions, they were calculated independently (SwissFEL 1 ps 21%, SwissFEL 10 ps 28%, SACLA 100 ps 22%)

### Refinement of light-activated states

The dark state model from SwissFEL was used as an initial model for refinement with the extrapolated data for the 1ps and 10 ps time points in *Phenix.refine*^67^, interactive model building with *Coot*^68^ was performed to fit the model to the 2*F_extra_* - *F_calc_* maps and to remove water molecules lacking electron density. The same iterative procedure was performed starting with the SACLA 100 ps data, using the SACLA dark state model as a starting point.

### Residue numbering

In addition to a number according to their position in the sequence, residues in rhodopsin are also assigned a ‘general’ number according to the Ballesteros-Weinstein (BW) scheme^1^. The BW general number consists of two numbers separated by a dot, where the first denotes the helix (1 to 8) and the second the position relative to the most-conserved residue in that helix, arbitrarily assigned to 50. For instance, Glu113^<u>3.28</u>^ denotes that the counterion Glu113 is located in TM3 and twenty-two residues before the most conserved residue in TM3 (Arg135^<u>3.50</u>^).

### QM/MM calculations

The initial coordinates for geometry optimization were taken from the TR-SFX crystallographic structures reported in this work. The pKa values at pH 9.0 of titratable amino acid residues in the protein were obtained with the PROPKA program^71,72^. Subsequently, the program tleap from the AMBER software package was used to protonate the protein by considering the previously calculated pKa values^73^. The geometries were optimized using hybrid quantum mechanics/molecular mechanics (QM/MM)^74^. In the simplest system, the QM part consists only of the retinal chromophore and the sidechain of Lys296 that forms the PSB. The hydrogen link atom (HLA) scheme^75^ was used to place the QM/MM boundary in between the Cδ and Cε atoms of the Lys296 sidechain. We also considered two more extended QM regions that include a) the proximal counterion (Glu113^<u>3.28</u>^) and b) the proximal counterion (Glu113^<u>3.28</u>^), Tyr268^<u>6.51</u>^, and water WAT01. The QM part was described using the BP86-D3(BJ) functional^76,77^ in conjunction with the cc-pVDZ basis set^78^ and the def2/J auxiliary basis set for the resolution of identity^79^. The Chain of Spheres exchange (COSX) algorithm was used in combination with the resolution of identity for the Coulomb term (RI-J). The remaining proteins were treated with the Amber ff14SB force field^80^. The TIP3P model was used to describe the water molecules^81^. The QM/MM optimizations were performed by using the quantum chemistry program Orca 5.0.2^82^ interfaced with the DL_POLY module of the ChemShell 3.7.1 software package^83, 84^. The minimized ground state geometries were used to calculate the vertical excitation energies at the RI-CC2 level of theory^85^ with frozen core orbitals and the cc-pVTZ basis set in association with the corresponding auxiliary basis^78^. The RI-CC2 calculations were performed with the Turbomole 7.5.1 program package^86^. All calculations were performed using the supercomputing facilities at the Paul Scherrer Institute.

