## Supplementary Data for "ULTRAFAST STRUCTURAL CHANGES DIRECT THE FIRST MOLECULAR EVENTS OF VISION"

**Extended Data Fig. 1 | Rhodopsin molecules packing in the crystal lattice of space group P22$_1$2$_1$**

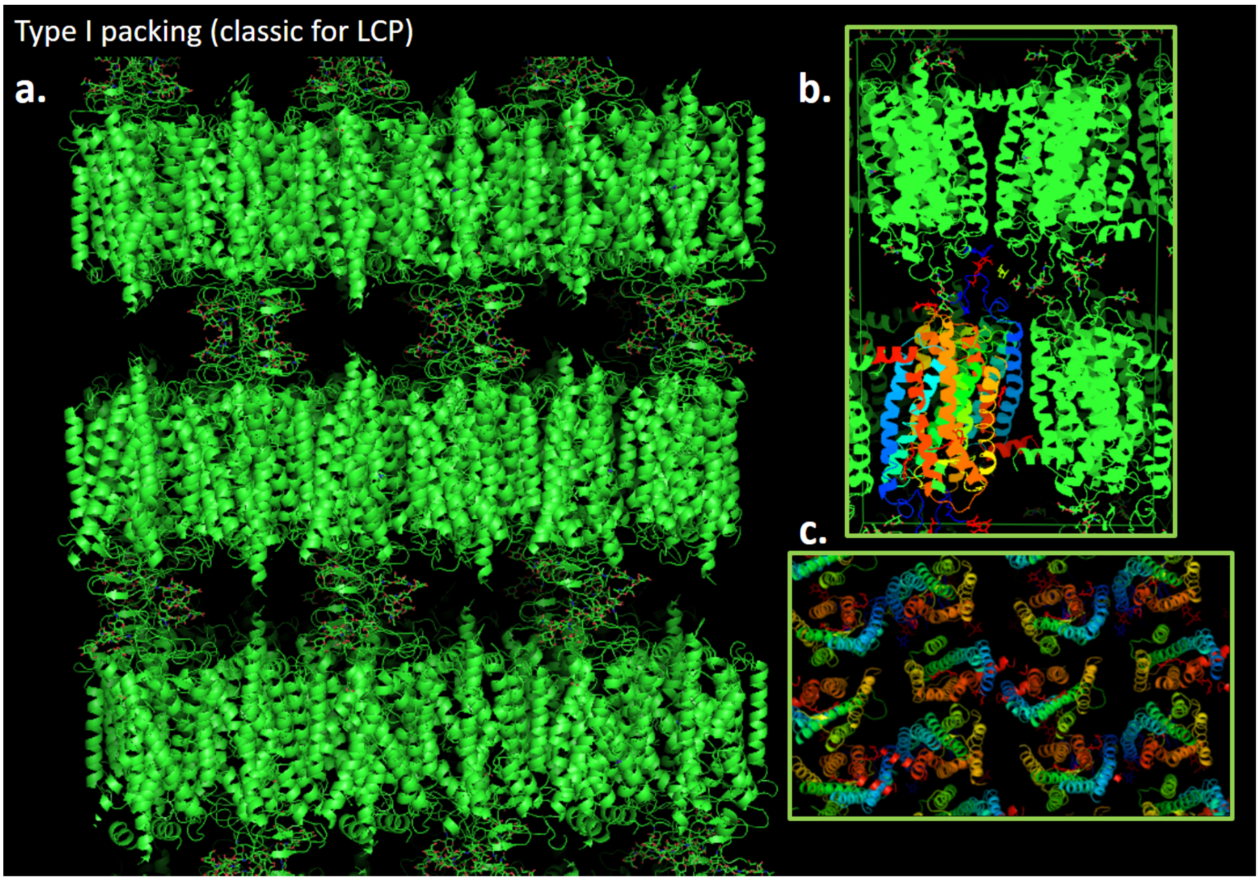

Bovine rhodopsin crystals obtained with the lipidic cubic phase method reveal a typical molecule packing of type I, consisting of a well-ordered stacking of 2D-crystals **(a.)** The 2D-crystals contact each other through the glycosyl groups of Asn2 and Asn15 of the rhodopsin N-termini, generating head-to-head crystal contacts in the c-dimension. **(b.)** View of the potential physiological dimer[28] contacting the transmembrane domains 1 (in blue) of each of the two rhodopsin molecules. **(c.)** top view of the molecules arrangement. The "Spectrum" rainbow colour transforms gradually from the TM1 in blue to the TM7 and the amphipathic helix 8 in red.

**Extended Data Fig. 2 | lattice translation correction**

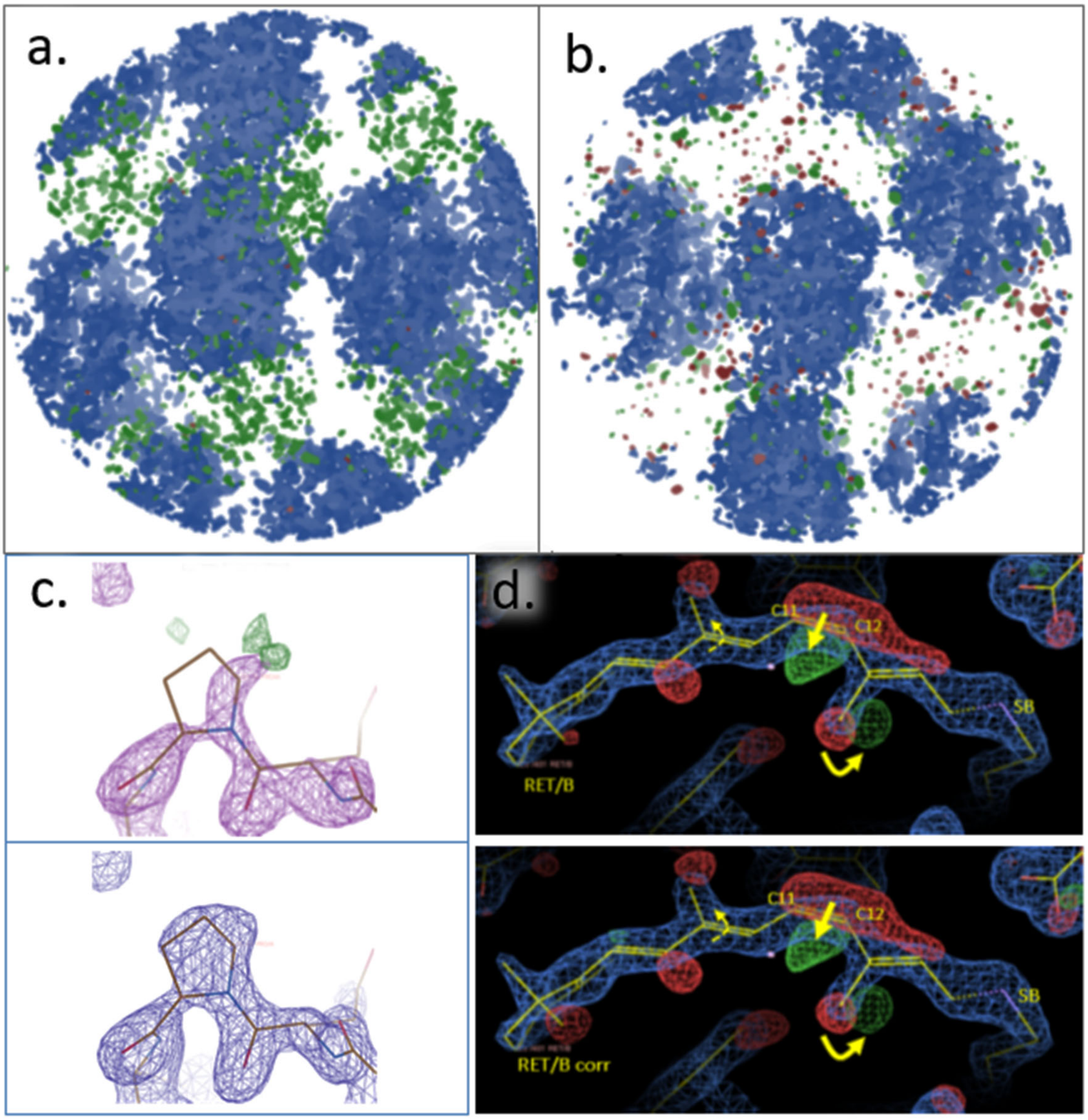

Despite a straight forward molecular replacement (Phaser MR, Phenix [67], see Methods) and a solution harbouring a $P22_12_1$ space group, the dark rhodopsin data analysis indicated the presence of more than one off-origin peaks in the Patterson function. In particular, the Patterson peak at td = (0,0.245,0)(SwissFEL) and (0,0.243,0)(SACLA) were attributed to the presence of two translation-related domains within the crystals. Accordingly, a ghost density was identified (**a**) in the Fo-Fc map (in green and red, contoured at 3.83 rmsd) partially overlapping with the 2Fo-Fc map (blue, 2.7 rmsd) from which the rhodopsin model was built in. **b)** After correction of the single domain X-ray intensities according to Wang et al [65] (see **Methods**), the overall rhodopsin electron density map 2Fo-Fc displays less ghost electron density (green and red). More locally at a few affected amino acid residues location, e.g. compare the electron density of Pro34/A in the lower panel of **(c)** (corrected) to upper (original). **d)** Importantly, the retinal binding pocket of rhodopsin was not affected by the density overlapping and correction, and the density after correction (lower panel) shows only minor changes in the difference map compared to the original data (upper panel).

**Extended Data Fig. 3 | Time resolved serial femtosecond crystallography**

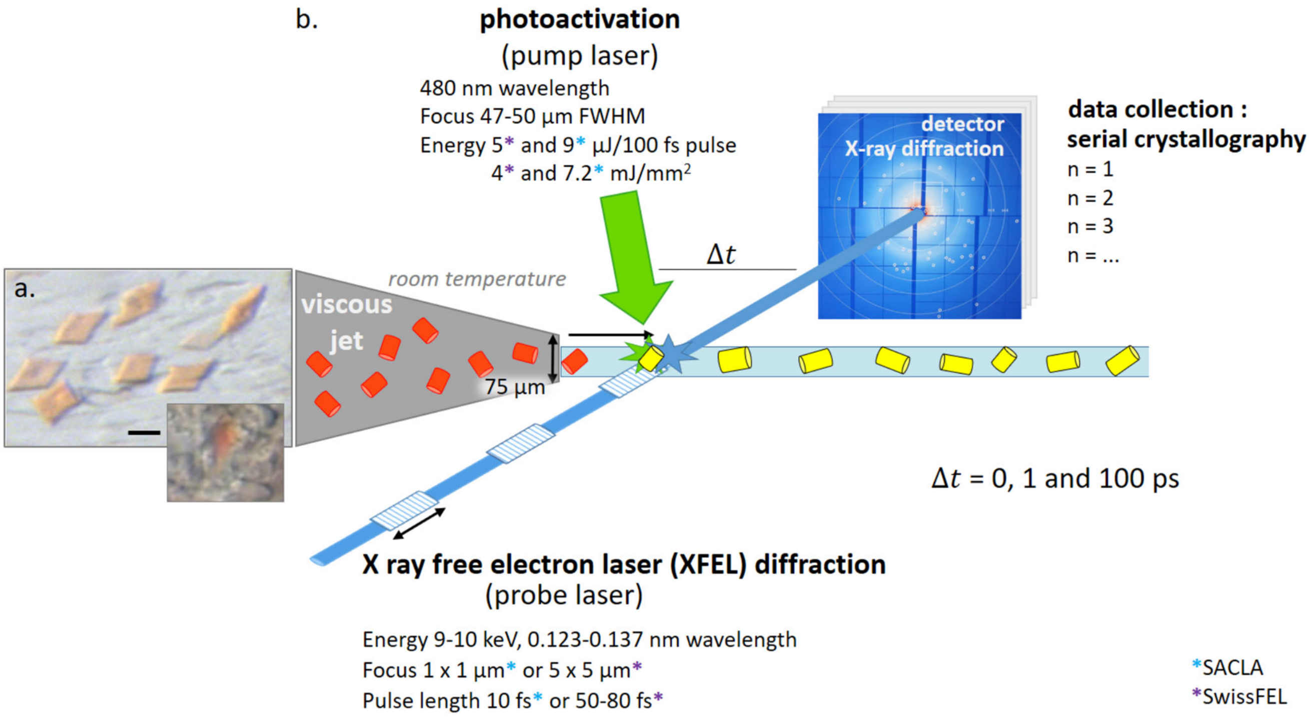

Time resolved serial femtosecond crystallography (TR-SFX)[87] was conducted using an X-ray free electron laser. Crystal plates (20 µm large and about 1.5 µm thick) made of rhodopsin purified from bovine retinae (**a. panel**, scale bar is 20 µm) were subjected to a pump and probe experiment (**b. panel**) triggered by a photoactivation at 480nm (pump laser). Briefly, rhodopsin crystals grown in lipidic cubic phase (LCP) in the darkness were successively injected (viscous jet) in the light of a pump laser and probed for X-ray diffraction after various time-delays ($\Delta t$ from 1 to 100 picoseconds) using the X-ray free electron laser from SACLA (Japan)(blue asterisk) or SwissFEL (Switzerland)(black asterisk) under the regime "diffract-before-destroy"[88]. The processing was done both on the fly and home using the CrystFEL software[62] combining 30'000 of the diffraction patterns generated by each crystal into a dataset. The inset of the **a. panel** illustrates the opacity of the lipidic phase around the crystal when the rhodopsin crystals are produced in large quantity, hampering spectroscopic experiments on the TR-SFX sample.

**Extended Data Fig. 4 | Photon absorption of rhodopsin sample analysed by TR-XSS**

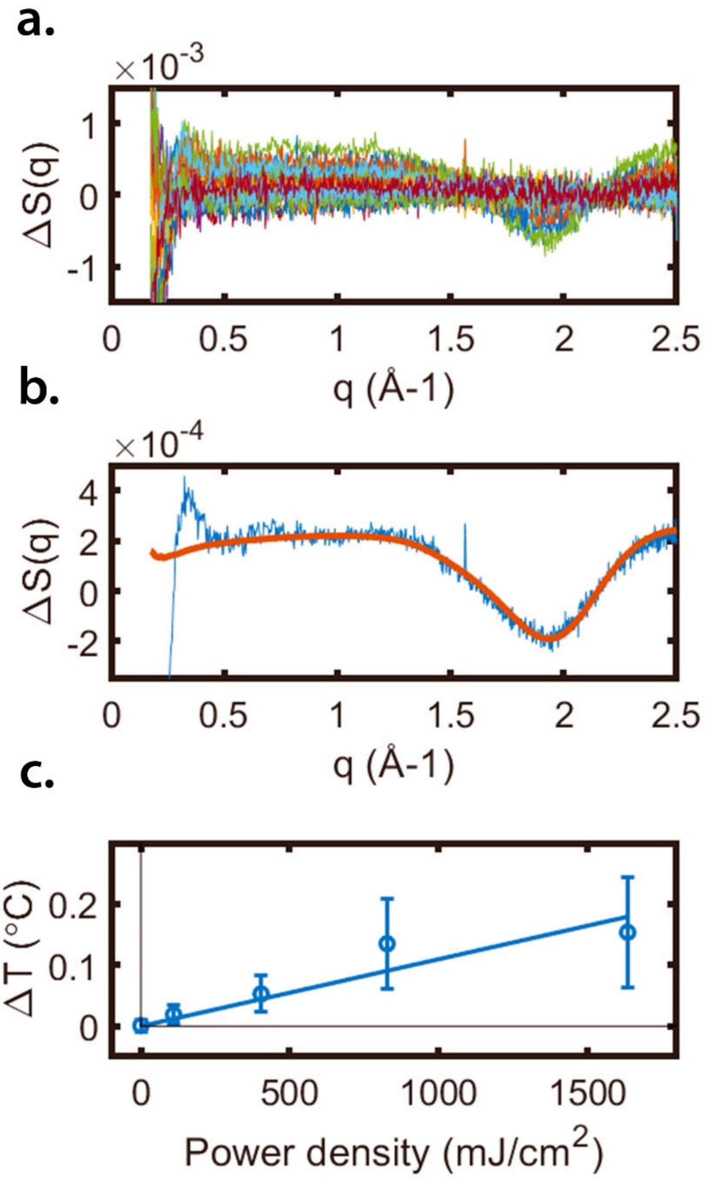

Time-resolved X-ray solution scattering (TR-XSS) studies of visual rhodopsin using XFEL radiation. **a**) TR-XSS difference data (laser on minus laser off) recorded at the LCLS from detergent solubilized samples of rhodopsin for the time-delays 10 ps ≤ Δt ≤ 1 µs at various laser power densities. **b**) Principal singular value decomposition (SVD) component (blue line) from samples of visual rhodopsin indicating laser induced heating (characteristic curve from 0.5 $Å^{-1} \leq q \leq$ 2.5 $Å^{-1}$) as well as oscillations usually associated with protein induced structural changes (visible from 0.25 $Å^{-1} \leq q \leq$ 1.0 $Å^{-1}$). An experimental difference X-ray scattering curve due to heating alone (red line) recorded from detergent solubilized samples of a photosynthetic reaction centre using synchrotron radiation [55], is used to calibrate this laser induced heating. **c**) Laser induced heating in samples of visual rhodopsin measured by TR-XSS when using a 480 nm fs laser pulse with a fluence of 110 mJ/cm$^2$; 400 mJ/cm$^2$; 830 mJ/cm$^2$; and 1640 mJ/cm$^2$, where the fluence is averaged across the FWHM of the laser spot. Negative time-delays were used as a control (plotted at zero fluence). Uncertainty bars are estimated from the standard deviation of a sequence of measurements divided by the square root of the number of independent observations.

**Extended Data Fig. 5 | Anisotropic breathing motion of rhodopsin.**

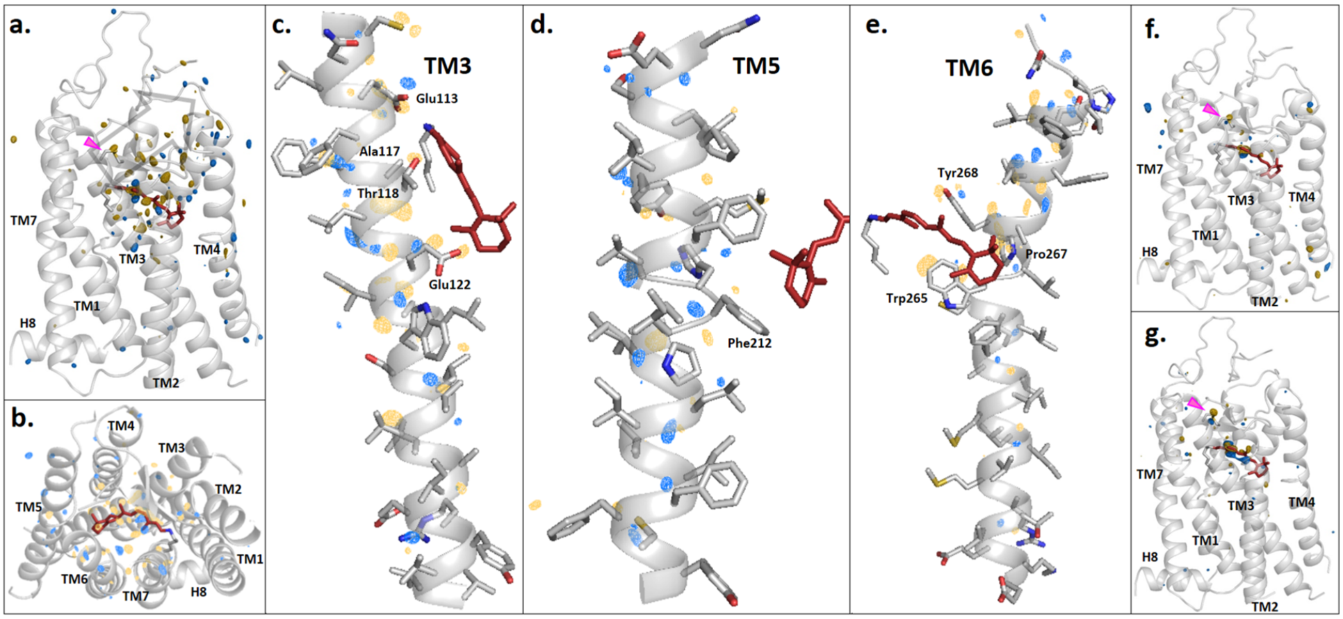

Comparison of the overall conformational changes in rhodopsin photoactivated for 1, 10 and 100 picoseconds. The difference electron density map (Fobs(1ps-light)-Fobs(dark) contoured at 4.2 rmsd) from the dataset of 1ps-illuminated rhodopsin superimposed on the rhodopsin dark state structure model (grey) (**a-b**) shows strong signals (blue=positive density; yellow=negative density) on the retinal molecule (red) demonstrating the early isomerisation. Surrounding the retinal, other changes occur at the amino acids level and in a directed way, towards the extracellular side (grey arrow of **panel a**), along TM5 and TM6 (**panel b**). This anisotropic breathing motion can be detected in the upper part of TM3 (**c**), TM5 (**d**) and TM6 (**e**). After 10 ps (**panel f**) and 100 ps (**panel g**) of photoactivation, most of the conformational concerted motion changes are reset (**f, g and Table 3**), and a few amino acids only will not revert –like the disulfide bridge Cys110-Cys187 (pink arrow)- and take part to further changes along the photoactivation pathway (**Table 3**).

**Extended Data Fig. 6 | Conformation of retinal after 1, 10 and 100 ps of rhodopsin photoactivation using TR-SFX.**

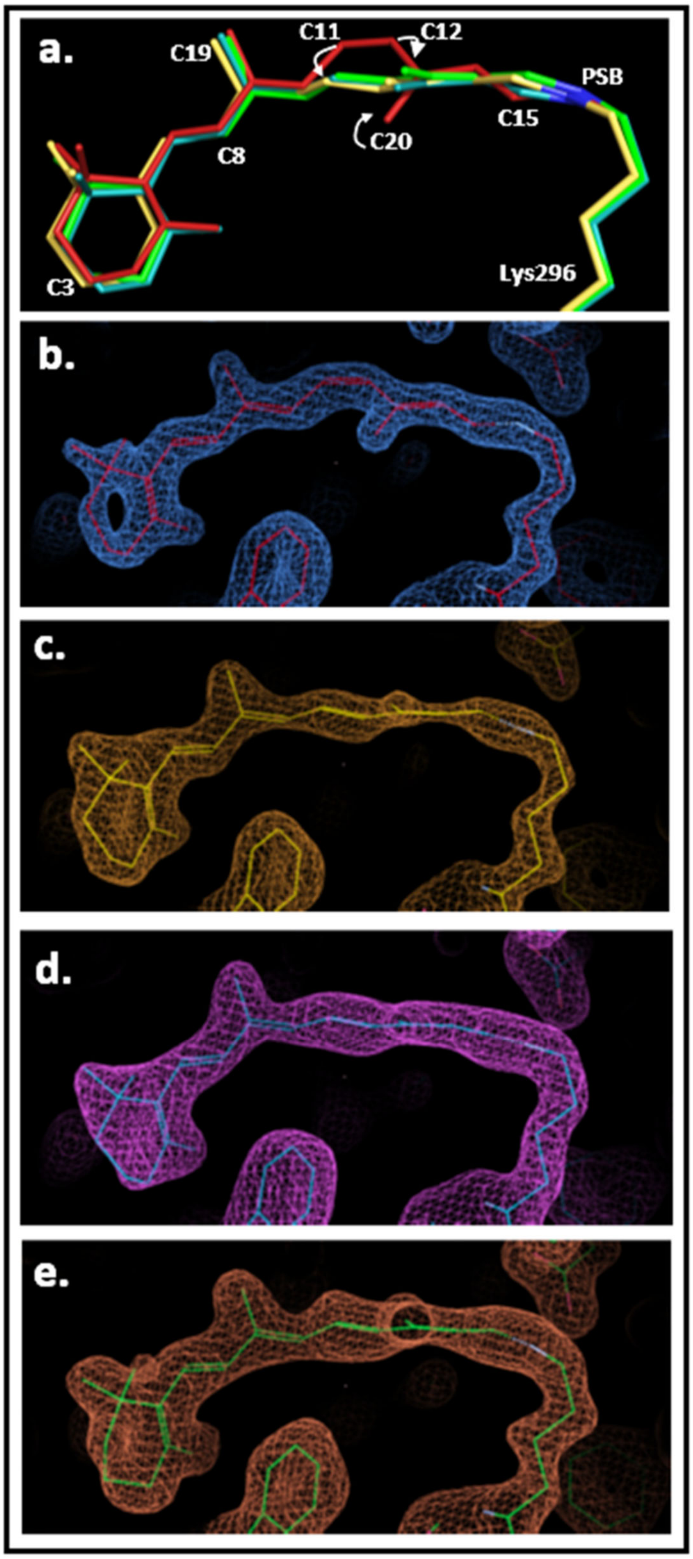

Retinal conformational changes until 100 ps. **a)** The superimposition of the retinal TR-SFX models in the dark and 1 to 100 ps photoactivation time delays highlights the main differences: the cis-to-trans isomerisation at C11-C12 and th concomitant rotation of the C20-methyl at C13. Beside a slight tilt of the β-ionone ring, the difference between 1, 10 and 100 ps-structures is a slight relaxation of the polyene chain towards planarity. **b)** Original electron density map around the retinal in the rhodopsin dark state obtained by SFX (2Fo-Fc map contoured at 2.5 rmsd) and the resulting refined model in red. **c)** Extrapolated electron density map around the retinal of 1ps-photoactivated rhodopsin obtained by TR-SFX (2Fext-Fc map contoured at 1.9 rmsd) and the resulting refined model in yellow. **d)** Extrapolated electron density map around the retinal of 10ps-photoactivated rhodopsin obtained by TR-SFX (2Fext-Fc map contoured at 1.9 rmsd) and the resulting refined model in blue. **e)** Extrapolated electron density map around the retinal of 100ps-photoactivated rhodopsin obtained by TR-SFX (2Fext-Fc map contoured at 1.9 rmsd) and the resulting refined model in green.

**Extended Data Fig. 7 | Schiff base – counterion Glu113 and neighbouring water hydrogen bond network after 1 picosecond of photoactivation.**

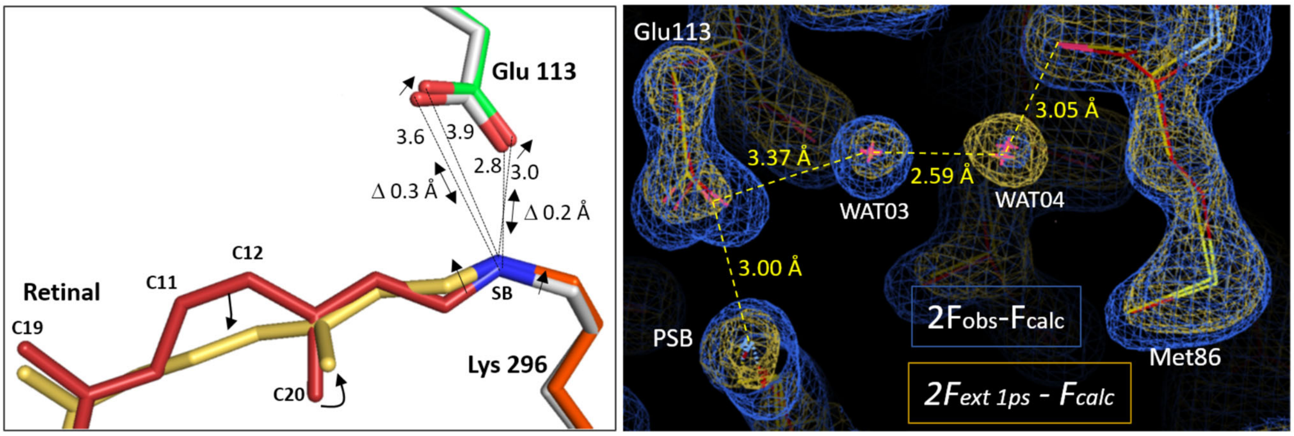

**Left panel**: influence of the C11-C12 isomerisation on the Schiff base conformation and distance to counterion. The two models of the rhodopsin are superimposed. The retinal after 1 ps of photoactivation (yellow (orange Lys296)) is showing an all-trans conformation and the C15 of the C14-C15-NZ plane at the SB displays a slight kick towards the extracellular space compared to the structure of the dark state (red). The counterion Glu113$^{3.28}$ moves accordingly, in the same direction of about 0.2-0.3Å. **Right panel**: from the two water molecules WAT03 and WAT04 forming a bridge between the counterion Glu113$^{3.28}$ and Met86$^{2.53}$ and contacting Ala117, Phe91 and Phe116, WAT04 has gained order. The two rhodopsin molecules models (dark in red; 1 ps in yellow) are contoured with their respective electron density maps, in blue (2Fo-Fc contoured at 1.3 rmsd) and in orange (2Fextrapolated-Fc contoured at 1.3 rmsd). By $\Delta t$ = 100 ps we observe a reset of the occupancy, which is similar to that of the dark state structure.

Water nomenclature: WAT 01 (chain C / HOH #01) at the tip of C20/RET; WAT 02 (chain C / HOH #02) at Ser 186; WAT 03 (chain C / HOH #103) proximal to counterion Glu113 (Gly90, Phe91, Ala 117); WAT 04 (chain C / HOH #119) at Met86 (Phe91, Phe116, Ala 117), low occupancy increasing by $\Delta t$ = 1 ps, reseting by $\Delta t$ = 100 ps.

**Extended Data Fig. 8 | Interactions of retinal with its binding pocket at 0, 1 and 100 ps of photoactivation.**

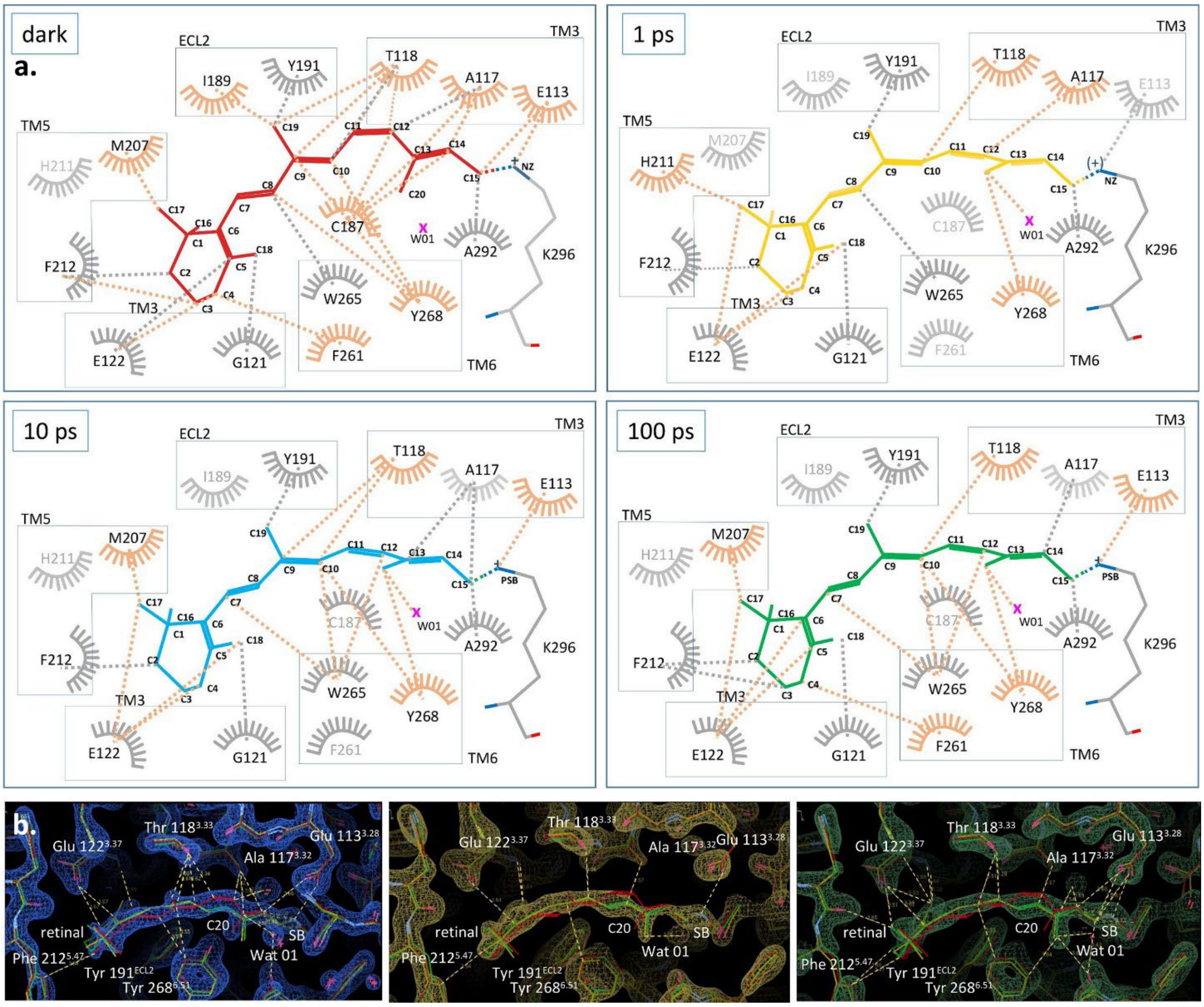

**Upper panels: LIGPLOT[90] view of retinal interactions within the rhodopsin binding site** (set distance < 3.6 Å) at different time-delays of photoactivation, dark state, 1, 10 and 100 ps-photoactivated states. The amino acids and dashed lines of interatomic interactions labelled in orange are the site of major changes, showing new interactions, e.g. with water W01 or losing contact with Cys 187.

**Lower panels: "residues environment distances" from retinal measured by the *Coot* software[68].** The two rhodopsin models after 1 ps (yellow) and 100 ps (green) of photoactivation were superimposed using Secondary Structure Matching with the rhodopsin dark state model (red) and the residues environment distances were drawn with a cut off at 3.7 Å using dashed lines from the retinal in the dark (**left panel**), 1 ps (**middle panel**) and 100 ps (**right panel**).

**Extended Data Table 1 | Crystallographic data for rhodopsin structures after 1 picosecond- and 100 picoseconds of photoactivation, compared to the dark state.**

Dark states collected at the SwissFEL and SACLA (SFX, laser off):

| | **Dark state SwissFEL** (Combined SwissFEL 2018 + 2020) | **Dark state SACLA** |
|---|---|---|
| Resolution Range (Å) | 16.10-1.80 (1.86-1.80) | 10.47-1.80 (1.86-1.80) |
| Unit Cell | a=61.51 Å, b=91.01 Å, c=151.11 Å, α=90.0°, β=90.0°, γ=90.0° | a=61.29 Å, b=90.81 Å, c=150.51 Å, α=90.0°, β=90.0°, γ=90.0° |
| Space group | $P\ 2\ 2_1\ 2_1$ | $P\ 2\ 2_1\ 2_1$ |
| Measured reflections | 65,870,940 (4,371,475) | 105,018,485 (7,436,654) |
| Unique reflections | 79,305 (7,852) | 78,209 (7,715) |
| Multiplicity | 830.6 (556.7) | 1,342.8 (963.9) |
| Completeness | 100.0% (100.0%) | 100.0% (100.0%) |
| I/σI | 7.62 (0.95) | 8.87 (1.32) |
| $R_{split}$ (%) | 8.21 (109.06) | 6.75 % (77.70 %) |
| CC* | 0.9982 (0.8947) | 0.9994 (0.9219) |
| $CC_{1/2}$ | 0.9926 (0.6672) | 0.9977 (0.7391) |
| Translation vector (Td) | 0.245 | 0.243 |
| Translated fraction (*k*) | 0.22 | 0.13 |
| **Refinement** | | |
| Resolution (Å) | 16.10 - 1.80 | 10.47 - 1.80 |
| $R_{work}/R_{free}$ | 21.46 / 24.74 | 19.79 / 22.32 |
| *No. of Atoms* | | |
| Protein | 4971 | 4970 |
| Ligand | 526 | 539 |
| Water | 174 | 168 |
| *B-factors* | | |
| Protein (Å$^2$) | 32.2 | 30.7 |
| Ligand (Å$^2$) | 48.2 | 48.4 |
| Water (Å$^2$) | 39.9 | 39.0 |
| *Ramachandran* | | |
| Favoured | 96.81% | 97.14% |
| Allowed | 3.19% | 2.86% |
| Outliers | 0.00% | 0.00% |
| *R.M.S.D deviations* | | |
| Bond lengths (Å) | 0.01 | 0.01 |
| Bond angles (°) | 0.91 | 0.96 |

**Extended Data Table 1 |** Crystallographic data for 1, 10 and 100 ps-photoactivated rhodopsin (TR-SFX)

| | **1 ps SwissFEL** | **10 ps SwissFEL** | **100 ps SACLA** |
|---|---|---|---|
| Resolution Range (Å) | 16.10-1.80 (1.86-1.80) | 16.10-1.80 (1.86-1.80) | 10.47-1.80 (1.86-1.80) |
| Unit Cell | a=61.51 Å, b=91.01 Å, c=151.11 Å, α=90.0°, β=90.0°, γ=90.0° | a=61.51 Å, b=91.01 Å, c=151.11 Å, α=90.0°, β=90.0°, γ=90.0° | a=61.29 Å, b=90.81 Å, c=150.51 Å, α=90.0°, β=90.0°, γ=90.0° |
| Space group | $P\ 2\ 2_1\ 2_1$ | $P\ 2\ 2_1\ 2_1$ | $P\ 2\ 2_1\ 2_1$ |
| Measured reflections | 23,497,455 (1,454,025) | 47,097,450 (3,363,550) | 53,114,217 (3,761,378) |
| Unique reflections | 79,304 (7,852) | 79,031 (7,852) | 78,209 (7,715) |
| Multiplicity | 296.3 (185.2) | 595.9 (428.4) | 679.1 (487.5) |
| Completeness | 100.0 % (100.0 %) | 99.7 % (100.0%) | 100.0% (100.0%) |
| I/σI | 5.76 (0.79) | 5.70 (0.72) | 6.20 (0.94) |
| $R_{split}$ (%) | 11.45 (124.17) | 10.24 % (138.78 %) | 9.68 % (111.03 %) |
| CC* | 0.9934 (0.8759) | 0.9979 (0.8209) | 0.9987 (0.8519) |
| $CC_{1/2}$ | 0.9741 (0.6224) | 0.9915 (0.5081) | 0.9947 (0.5695) |
| Translation vector (Td) | 0.246 | 0.245 | 0.243 |
| Translated fraction (*k*) | 0.22 | 0.21 | 0.13 |
| **Refinement** | | | |
| Resolution (Å) | 9.99 - 1.80 | 9.99 - 1.80 | 10.47-1.80 |
| Activation Level (%) | 21% | 28% | 22 % |
| $R_{work}/R_{free}$ | 35.5 / 40.1 | 30.8 / 34.7 | 33.0 / 37.1 |
| *No. of Atoms* | | | |
| Protein | 4971 | 4971 | 4966 |
| Ligand | 516 | 467 | 511 |
| Water | 164 | 115 | 140 |
| *B-factors* | | | |
| Protein (Å$^2$) | 27.67 | 45.53 | 30.81 |
| Ligand (Å$^2$) | 40.03 | 58.94 | 45.69 |
| Water (Å$^2$) | 31.78 | 47.75 | 33.16 |
| *Ramachandran* | | | |
| Favoured | 96.14 % | 96.14 % | 95.95 % |
| Allowed | 3.86 % | 3.86 % | 4.05 % |
| Outliers | 0.00 % | 0.00 % | 0.00 % |
| *R.M.S.D deviations* | | | |
| Bond lengths (Å) | 0.01 | 0.01 | 0.01 |
| Bond angles (°) | 0.860 | 0.832 | 0.735 |

**Extended Data Table 2 | Vertical excitation energies computed by quantum mechanics/molecular mechanics optimization.**

| Structure | exp. $\lambda_{max}$ nm (eV) | QM Region | $\Delta E_{calc}$ nm (eV) | Oscillator strength | $\Delta\Delta E_{calc}$ (1 ps - dark) nm | $\Delta\Delta E_{exp}$ (1 ps - dark) nm |
|---|---|---|---|---|---|---|
| Dark | 498 (2.49) | RET+K296 | 492 (2.52) | 1.55 | - | - |
| | | RET+K296+E113 | 456 (2.72) | 1.61 | - | - |
| | | RET+K296+E113+Y268+W001 | 464 (2.67) | 1.32 | - | - |
| 1 ps | 529 (2.34) | RET+K296 | 522 (2.37) | 1.81 | 30 | 31 |
| | | RET+K296+E113 | 490 (2.53) | 1.87 | 34 | |
| | | RET+K296+E113+Y268+W001 | 495 (2.51) | 1.75 | 31 | |

Vertical excitation energies (ΔEcalc) and oscillator strengths for dark and 1ps photoactivated rhodopsin. Three QM regions were considered: the basic retinal/Schiff base system (RET+K296), plus the counterion (RET+K296+E113), and plus Tyr268$^{6.51}$ and a nearby water molecule (RET+K296+E113+Y268+W001). The values were computed with the RI-CC(2) method using the cc-pVTZ basis set. The experimental values for the absorption maxima of each state (exp. $\lambda_{max}$) are provided for comparison. We observe that the computed ΔEcalc values for the simplest QM region considered (retinal + Schiff base, highlighted in grey) agree well with the experimental absorption maxima and, more importantly, they reproduce very well the expected red shift in the transition to Batho-Rh (30 nm, compared to the experimental value of 31 nm). Making the QM region more complex results in a consistent ~30 nm blue shift in ΔEcalc for both dark and 1 ps rhodopsin. As a result, these shifts cancel out to still provide a very good estimate for the rhodopsin-to-Batho-Rh transition (34 and 31 nm, compared to the experimental value of 31 nm).

**Extended Data Table 3 | Amplitude of amino acids shifts (Å) after 1ps photoactivation.**

| Amino acids | 1 ps<br>Shift side chain // main chain in mol a | 1 ps<br>Shift side chain // main chain in mol b | 100 ps<br>same - reset - or larger |
|---|---|---|---|
| Lys 296 | 0.35 // 0.20 | 0.25 // 0.21 | reset |
| Glu 113 | 0.24 // 0.22 | 0.31 // 0.31 | same as by 1 ps |
| Wat02 (Glu 113) | 0, reduced occupancy | 0, reduced occupancy | |
| Wat03 (Met 86) | 0, higher occupancy | 0, higher occupancy | |
| Tyr 268 | 0.41 // 0.40 | 0.31 // 0.36 | reset |
| Wat 01 (C20) | 0.67 | 0.29 | same as by 1 ps |
| Tyr 191 | 0.38 // 0.23 | 0.38 // 0.23 | same as by 1 ps |
| Ala 117 | 0.28 // 0.31 | 0.20 // 0.30 | reset |
| Thr 118 | 0.26 // 0.33 | 0.33 // 0.35 | reset |
| Trp 265 | 0.32 // 0.27 | 0.30 // 0.29 | same as by 1 ps |
| Glu 181 | 0.23 // 0.0 | 0.39 // 0.18 | reset |
| Ser 186 | 0.33 // 0.20 | 0.35 // 0.24 | same as by 1 ps |
| Cys 187 | 0.30 // 0.24 | 0.27 // 0.18 | same as by 1 ps |
| Glu 122 | 0.28 // 0.23 | 0.18 // 0.19 | CA reset but not carbonyl |
| Phe 212 | 0.24 // 0.21 | 0.26 // 0.25 | reset |
| Met 207 | 0.33 // 0.22 | 0.12 // 0.13 | reset |
| Phe 261 | 0 // 0 | 0 // 0 | |
| Ala 269 | 0.36 | 0.33 | reset |
| Pro 267 | 0.38 | 0.30 | reset |
| Ala 292 | 0.35 | 0.47 | |
| Gly 121 | 0.2 | 0.22 | reset |
| Pro 215 | 0.17 | 0.19 | reset |
